# Intraspecific diversity in the mechanisms underlying abamectin resistance in a cosmopolitan pest

**DOI:** 10.1101/2022.11.25.517948

**Authors:** Ernesto Villacis-Perez, Wenxin Xue, Marilou Vandenhole, Berdien De Beer, Wannes Dermauw, Thomas Van Leeuwen

**Author notes:** These authors contributed equally to this work (Ernesto Villacis-Perez, Wenxin Xue). Corresponding author: Thomas Van Leeuwen postal address: Coupure links 653, 9000 Ghent.

## Abstract

Pesticide resistance relies on a myriad of mechanisms, ranging from single mutations to a complex and polygenic architecture, and it involves mechanisms such as target-site insensitivity, metabolic detoxification, or a combination of these, with either additive or synergistic effects. Several resistance mechanisms against abamectin, a macrocyclic lactone widely used in crop protection, have been reported in the cosmopolitan pest *Tetranychus urticae*. However, it has been shown that a single mechanism cannot account for the high levels of abamectin resistance found across different mite populations. Here, we used experimental evolution combined with bulked segregant analyses to map quantitative trait loci (QTL) associated with abamectin resistance in two genetically unrelated strains of *T. urticae*. In these two independent QTL mapping experiments, three and four QTLs were identified, of which three were shared between experiments. Shared QTLs contained genes encoding subunits of the glutamate-gated chloride channel (GluCl) and harboured previously reported mutations, including G314D in *GluCl1* and G326E in *GluCl3*, but also novel resistance candidate loci, including DNA helicases and chemosensory receptors. Surprisingly, the fourth QTL, present only in only one of the experiments and thus unique for one parental resistant strain, revealed a non-functional variant of *GluCl2*, suggesting gene knock-out as resistance mechanism. Our study uncovers the complex basis of abamectin resistance, and it highlights the intraspecific diversity of genetic mechanisms underlying resistance in a cosmopolitan pest.

## 1 INTRODUCTION

Insecticidal and acaricidal crop protection compounds that target conserved physiological processes, such as respiration or neural function, are widely used as control agents against agricultural pests. However, pests can develop resistance to these compounds, as observed in many arthropod species (Hawkins et al., 2018; Mota-Sanchez & Wise, 2022; Sparks et al., 2021; Van Leeuwen et al., 2010). Pesticide resistance is an evolutionary process with detrimental applied consequences, as it threatens efforts to build a sustainable agriculture framework (European Commission, 2019). Resistance can arise from changes in the coding sequence of the target proteins, i.e., toxicodynamic or target-site resistance; or from changes that affect the metabolism, excretion, transport or penetration of toxins, i.e., toxicokinetic resistance (Feyereisen et al., 2015; Van Leeuwen & Dermauw, 2016). In addition, a synergism between toxicodynamic and toxicokinetic mechanisms is likely to underlie resistance in particularly highly resistant pest populations (De Beer, Villacis-Perez, et al., 2022; Samantsidis et al., 2020; Zhang et al., 2021). Cases of target-site insensitivity underlying resistance, as well as metabolic detoxification and transport of pesticides via enzyme families such as cytochrome P450 mono-oxygenases (CYPs), glutathione S-transferases (GSTs), carboxyl-/cholinesterases (CCEs) and ATP-binding cassette (ABC) transporters, have been reported across different taxa (Feyereisen et al., 2015; Van Leeuwen & Dermauw, 2016). In addition, enzymes of the uridine diphosphate (UDP)-glycosyltransferases (UGTs) and transporters of the Major Facilitator Superfamily (MFS) have been associated with resistance (Ahn et al., 2012, 2014; De Beer, Vandenhole, et al., 2022; Dermauw et al., 2012; Snoeck, Pavlidi, et al., 2019). Finally, horizontally transferred genes from bacteria and fungi (Wybouw et al., 2016), such as the intra-diol ring cleavage dioxygenases (DOGs) from spider mites, can convey unexpected metabolic capacities that were previously overlooked in studying xenobiotic metabolism, such as aromatic ring cleavage (Njiru et al., 2022). Most often, increased expression of these enzymes and transporters is associated with resistance, but the underlying gene regulatory mechanisms have in most cases remained elusive. Recently, the quantification of allele-specific expression was used to uncover that *trans*-driven gene regulation is common amongst detoxifying gene families such as P450s and DOGs in several resistant strains of the polyphagous arthropod *Tetranychus urticae* (Kurlovs et al., 2022). Yet, despite ample evidence on the proximal physiological processes and enzymatic pathways related to mechanisms of pesticide resistance in arthropods, the evolutionary origins of the underlying traits, the genetic structure, and the specific genomic targets of pesticide selection, as well as evidence for the synergism between different resistance mechanisms, are far from completely elucidated (Fotoukkiaii et al., 2021; Hawkins et al., 2018; Neve et al., 2014; Singh et al., 2021; Van Leeuwen & Dermauw, 2016).

The genetic basis of pesticide resistance can be highly variable in arthropods, ranging from simple and monogenic, to complex and polygenic (Hemingway et al., 2004; Li et al., 2007; Van Leeuwen & Dermauw, 2016; Wybouw, Kosterlitz, et al., 2019). Thus, elucidating the genetic basis of resistance requires an unbiased approach that allows to map genomic loci with a high resolution. One such approach is an experimental evolution-based, population-level bulked segregant analysis (BSA), as recently developed for spider mites (see Kurlovs et al. (2019) for a review). Spider mites are particularly suitable for high-resolution genetic mapping approaches. Strains can be experimentally inbred and crossed in order to create segregating populations. Due to their short generation time and exponential population growth, segregating populations can be easily expanded to thousands of individuals in a relatively confined space, which promotes a high number of recombination events that can help resolve causal loci to very narrow genomic regions (Kurlovs et al., 2019). Indeed, recent research has shown that the spider mite *Tetranychus urticae* is an ideal organism to study the genetic basis of adaptive evolution, both in experimental and field settings (Belliure et al., 2010; Bryon et al., 2017; Fotoukkiaii et al., 2021; Kurlovs et al., 2019; Snoeck, Kurlovs, et al., 2019; Sugimoto et al., 2020; Van Leeuwen et al., 2012; Villacis-Perez et al., 2021; Wybouw, Kosterlitz, et al., 2019; Wybouw, Kurlovs, et al., 2019). *T. urticae* is a generalist herbivore that harbours high levels of intraspecific genetic variation, possibly associated to adaptation to different host plants (Villacis-Perez et al., 2021). Further, cases of resistance to nearly all classes of acaricides, pesticides to combat mites and ticks, have been reported for this species (Mota-Sanchez & Wise, 2022; Sparks & Nauen, 2015; Van Leeuwen et al., 2020; Van Leeuwen & Dermauw, 2016). Even though common genetic variants associated with resistance to different classes of compounds are repeatedly identified in field mite populations (Van Leeuwen et al., 2020), it is less clear whether alternative genetic variants, mechanisms, or different combinations of resistance factors underlie the phenotype of resistance across unrelated populations.

For example, abamectin, consisting of macrocyclic lactones avermectin B1a and avermectin B1b, has been used as an effective acaricide against *T. urticae* for the last thirty years, but cases of resistance have only been documented in the last decade (Dermauw et al., 2012; Kwon et al., 2010; Memarizadeh et al., 2013; Monteiro et al., 2015; Sato et al., 2005; Xu et al., 2018; Xue et al., 2020; Zhang et al., 2021). Multiple resistance mechanisms to abamectin have been described, including target-site insensitivity and oxidative metabolism, suggesting a complex genetic basis of abamectin resistance (Kwon et al., 2010; Mermans et al., 2017; Riga et al., 2014; Wang et al., 2017; Xue et al., 2020, 2021). Abamectin is an allosteric modulator that targets cys-loop ligand-gated ion channels in invertebrates, of which glutamate-gated chloride channels (GluCl) are the main target site in arthropods and nematodes (Clark et al., 1995; Dent et al., 2000; Ludmerer et al., 2002; Mermans et al., 2017; Sparks et al., 2021). Mutations in GluCl genes associated with resistance to avermectins have been identified in *Caenorhabditis elegans, Drosophila melanogaster, Plutella xylostella* and *T. urticae*, and in some cases these mutations have been functionally validated using different approaches, including two-electrode voltage-clamp electrophysiology, the creation of near-isogenic lines and classic backcrossing experiments with F2 screens (Choi et al., 2017; Dent et al., 2000; Dermauw et al., 2012; Ghosh et al., 2012; Hibbs & Gouaux, 2011; Kwon et al., 2010; Mermans et al., 2017; Riga et al., 2017; Wang et al., 2016, 2017; Xue et al., 2021). Enhanced oxidative metabolism of avermectins has also been reported as a resistance mechanism. Synergism experiments, gene-expression analysis, and dedicated assays with functionally expressed proteins have pointed towards the role of cytochrome P450s (P450s) in the detoxification of abamectin in *Leptinotarsa decemlineata*, *Bemisia tabaci, P. xylostella* and *T. urticae* (Ludmerer et al., 2002; Qian et al., 2008; Riga et al., 2014; Wang & Wu, 2007; Xue et al., 2020; Yoon et al., 2002; Zuo et al., 2021). Other resistance mechanisms, including sequestration and metabolic detoxification via UGTs and GSTs may also play a role in avermectin resistance in arthropods and nematodes (Ghosh et al., 2012; Pavlidi et al., 2015; Snoeck, Pavlidi, et al., 2019; Wang et al., 2018; Xue et al., 2020). So far, studies focusing on abamectin resistance have been mostly limited to investigating the role of individual candidate mechanisms and their associated genes. However, there is a clear need for a population genomic approach that points without bias to genomic loci (QTL) involved in resistance. This will lead to a better understanding of the genetic basis and the diversity of mechanisms associated with abamectin resistance.

In this study, we aim to identify the genetic basis of abamectin resistance. To do so, we used experimental evolution in combination with a population-level based bulked segregant analysis (BSA) and with next-generation sequencing to map quantitative trait loci (QTL) associated with abamectin resistance in two genetically unrelated strains of *T. urticae* (Kurlovs et al., 2019). Our results reveal how unrelated populations of a single arthropod species respond to abamectin selection, the (polygenic) structure of abamectin resistance, and the genes and mechanisms likely to underlie abamectin resistance.

## 2 MATERIALS AND METHODS

### 2.1 Plants and acaricide

Common bean (*Phaseolus vulgaris* cv. ‘Speedy’ or ‘Prelude’) plants were grown from seeds two weeks prior to experiments at 25 °C, 60% RH and 16:8 L:D photoperiod (hereafter referred to as ‘standard conditions’) inside a greenhouse. A commercial formulation of abamectin (Vertimec, 18 g/L suspension concentrate) was used for all assays.

### 2.2 *Tetranychus urticae* lines

Two abamectin-resistant populations, MAR-AB and ROS-IT (referred to as IT2 in Xue et al., 2020) and two abamectin-susceptible populations, SOL-BE and JP-RR were inbred in the laboratory, yielding four inbred lines previously described in Kurlovs et al., (2022): MAR-ABi from MAR-AB, SOL-BEi from SOL-BE, JP-RRi from JP-RR, and ROS-ITi from ROS-IT. The inbred lines were maintained at standard conditions in the laboratory on detached bean leaves resting on wet cotton wool to prevent cross-contamination. To assess the concentration of abamectin that is lethal to half of the mite population (LC_50_), dose-response curves were constructed as described previously (Fotoukkiaii et al., 2019; Van Leeuwen et al., 2004; Xue et al., 2020). For each line, LC_50_ values, slopes and 95% confidence limits were estimated using PoloPlus (LeOra Software, Berkeley, CA, USA, 2006).

### 2.3 Differential gene expression analysis between susceptible and resistant parental lines

A differential gene expression analysis between pairs of abamectin-resistant and susceptible parental lines, i.e. MAR-ABi versus SOL-BEi and ROS-ITi versus JP-RRi, was performed using previously available data described in Kurlovs et al., (2022). Briefly, in Kurlovs et al., 2022, total RNA was extracted from a pool of 100–120 adult females per line using the RNeasy plus mini kit (Qiagen, Belgium) with 4 replicates for strains MAR-ABi and JP-RRi and 5 replicates for strains ROS-ITi and SOL-BEi. The concentration and purity of the RNA samples were assessed using a DeNovix DS-11 spectrophotometer (DeNovix, Willmington, DE, USA) and checked visually via gel electrophoresis (1% agarose gel; 30 min; 100V). Illumina libraries were constructed using the Illumina Truseq stranded mRNA library prep kit and sequenced on the Illumina Hiseq3000 platform (PE150 bp) at NXTGNT (Ghent, Belgium). RNA reads were mapped using GSNAP (version 2018-07-04) based on the known splice sites in the *T. urticae* GFF annotation (Wybouw, Kosterlitz, et al., 2019) whilst enabling novel splice-site discovery (“--localsplicedist=50000 --novelend-splicedist=50000 --pairmax-rna=50000”). Reads aligning to coding sequences were counted using HTSeq (version 0.11.2) and default settings for reverse-stranded samples (“-s reverse”). The resulting read counts were used as input for the ‘DESeq2’ (version 1.36.0) package in R to determine differentially expressed genes (DEGs) between each pair of resistant and susceptible parental lines, using an absolute log_2_ fold change (Log_2_FC) ≥ 2 and a Benjamini-Hochberg adjusted p-value (padj) < 0.05).

A *de novo* transcriptome was assembled for MAR-ABi and SOL-BEi using approximately 30 million paired-end RNA reads of each line (one replicate from MAR-ABi and two replicates from SOL-BEi) and the Trinity v2.1.1 software (Grabherr et al., 2011) with default settings and the “-trimmomatic” option to remove adapter sequences.

### 2.4 Experimental evolution set-up

To identify the genomic responses to abamectin selection, two independent BSA experiments were conducted, referred to as aBSA and gBSA, at the University of Amsterdam and at the University of Ghent, respectively. The starting mapping population for aBSA was created by crossing 60 virgin females of the susceptible line SOL-BEi to 20 adult males of the resistant line MAR-ABi. The mapping population for gBSA was created by crossing 41 virgin females of the susceptible JP-RRi line to 21 adult males of the resistant ROS-ITi line. To quantify resistance in F1 hybrids, LC_50_ values, slopes and 95% confidence limits were estimated in females of each F1 hybrid population in a similar set-up as the parental lines. To quantify the degree of dominance of the resistant trait we used the formula of Stone (1968): *D* = (*2logX_2_ - logX_1_ - logX_3_*)/(*logX_1_ - logX_3_*), where *X_1_* is the LC_50_ value of the resistant parent, *X_2_* is the LC_50_ value of the F1 progeny, and *X_3_* is the LC_50_ value of the susceptible parent.

Approximately 200-300 F1 female individuals from each cross were used to infest 4-8 potted bean plants, placed inside climate cabinets (Panasonic MLR-352H-PE, Kadoma, Japan) at standard conditions. After four to five generations of population expansion (generation time is around 12 days under these conditions for *T. urticae*), approximately 350 females from each mapping population were used to infest the 10 control replicates, each consisting of a fresh potted bean plant placed inside a mite-proof cage surrounded by a soapy water barrier, in a greenhouse at standard conditions. Control replicates were expanded for two generations, with a constant supply of fresh bean plants. Ten treatment replicates for each BSA assay were created by spraying potted bean plants with abamectin at a starting concentration of 0.5 mg a.i. /L for aBSA and 1 mg a.i. /L for gBSA, until runoff. Sprayed plants were placed inside new mite-proof cages. Then, ~500 individuals from each control replicate were used to infest each of the treatment replicates to create a paired setup. Thus, control replicate 1 from aBSA was used to infest abamectin replicate 1 from aBSA, control replicate 1 from gBSA infested abamectin replicate 1 from gBSA, and so on, for a total of 10 abamectin treatment replicates per experiment (gBSA and aBSA). Abamectin concentrations were progressively increased over time throughout the experimental evolution experiments, allowing the populations to build up large numbers before increasing selection strength. Acaricide concentrations for the final rounds of selection were 20 mg a.i. /L for aBSA and 2 mg a.i. /L for gBSA. Unsprayed controls on bean were refreshed with potted plants when the spider mite populations reached high numbers, but before the plant was completely eaten by the mites. The total duration of the experiments was 9 and 5 months for aBSA and gBSA, respectively.

### 2.5 Abamectin susceptibility assays

To evaluate the effectiveness of abamectin selection, the survival of the paired control and abamectin-selected replicates was quantified with toxicity tests. Before performing the toxicity tests, populations were expanded on unsprayed bean plants for at least one generation to avoid possible maternal effects and to exclude the effect of acaricide preexposure in the treatment replicates. Between 20 and 30 adult females from each replicate were transferred to a 9cm^2^ square bean leaf disc placed on wet cotton cool, and subsequently sprayed with 1 ml of fluid at 1 bar pressure with a Potter spray tower for replicates of aBSA, or with 0.8 ml of fluid at 1 bar pressure with a Cornelis spray tower for replicates of gBSA, to obtain a homogeneous spray film (deposit of approximately 1.9 mg/cm^2^). Four leaf discs were sprayed for each abamectin concentration for each BSA replicate, and four discs were sprayed with water as a control. Unselected control replicates and abamectin-selected replicates were sprayed with 15 mg a.i. /L abamectin for aBSA and with 10 mg a.i. /L for gBSA. After spraying, leaf discs were kept in a climate chamber at standard conditions. After 24 hours, survival was determined by assessing if mites could walk normally after being prodded with a camel’s hair brush. The survival percentage of each disc was corrected by the averaged mortality of the water-sprayed controls using Schneider-Orelli’s formula (Puntener & Ciba-Geigy, 1981). The overall difference in the percentage of corrected survival between acaricide-selected treatments and unselected controls was analysed using a linear mixed effect model for each experiment in R (package ‘lme4’, version 1.1-26 (Bates et al., 2015)), with selection treatment as fixed factor and paired replicate as random factor.

### 2.6 DNA extraction

Genomic DNA (gDNA) was extracted from adult female mites from the four parental inbred lines and from each of the BSA replicates according to Van Leeuwen et al. (2008). Individuals were collected in Eppendorf tubes, flash frozen in liquid nitrogen and stored at −80 °C until gDNA extraction. Two tubes of approximately 400 adult females each were collected from every replicate. Individuals in each tube were homogenized with a mix of 782 μl of SDS buffer (2% SDS, 200 mM Tris-HCl, 400 mM NaCl, 10 mM EDTA, pH = 8.33), 3 μl RNase A (20-40 mg/ml) (Thermo Fisher Scientific, Waltham, MA, USA) and 15 μl proteinase K (10 mg/ml) (Sigma-Aldrich, St-Louis, MO, USA), followed by DNA extraction using a previously described phenol-chloroform-based protocol. Prior to adding isopropanol, the extracts from the two tubes were pooled and precipitated together to obtain sufficient DNA per population. gDNA quality and quantity of the samples were quantified using an ND-1000 NanoDrop (Thermo Fisher Scientific, Waltham, MA, USA) or a Denovix DS-11 spectrophotometer (DeNovix, Wilmington, DE, USA) and by running an aliquot on a 2% agarose gel electrophoresis (30 min, 100V).

### 2.7 Whole-genome sequencing, variant calling and quality control on predicted variants

For ROS-ITi, Illumina libraries were constructed at NXTGNT (Ghent University) using the NEBNext Ultra II DNA library prep kit and sequenced using an Illumina HiSeq 3000 platform, generating paired-end reads of 150 bp. For the other parental strains, SOL-BEi, MAR-ABi and JP-RRi, Illumina libraries were constructed with the Truseq Nano DNA library prep kit and sequenced using the Illumina HiSeq2500 platform at the Huntsman Cancer Institute of the University of Utah (Salt Lake City, UT, USA), generating paired-end reads of 125 bp. Genomic sequence reads of the four parental strains were publicly available in the NCBI Sequence Read Archive (SRA) under BioProject PRJNA799176 (Kurlovs et al., 2022). For all the experimental BSA replicates, Illumina libraries were constructed using the TruSeq Nano DNA sample preparation kit by Fasteris (https://www.fasteris.com/dna) and sequenced on an Illumina Novaseq 6000 Sequel platform, generating paired-end reads of 100 bp. Genomic sequence reads of all BSA replicates were deposited to the NCBI Sequence Read Archive under BioProject XXXXXX. Variant calling was performed as described in Snoeck, Kurlovs, et al. (2019). Briefly, reads were aligned to the reference Sanger draft *T. urticae* genome obtained from the London strain (Grbiċ et al., 2011) using the default settings of the BurrowsWheeler Aligner (version 0.7.17-r1188) (Li & Durbin, 2009), and then processed into position-sorted BAM files using SAMtools 1.11 (Li & Durbin, 2009) and the three pseudochromosome assembly of *T. urticae* (Wybouw, Kosterlitz, et al., 2019). Duplicates were marked using Picard tools (version 2.20.4-SNAPSHOT) (https://broadinstitute.github.io/picard). Joint variants were called across the 40 experimental selected and unselected populations and the four parental lines using GATK’s (version 4.1.7.0) (McKenna et al., 2010). The GenotypeGVCFs tool of GATK was used to produce a variant call format (VCF) file containing single nucleotide polymorphisms (SNP’s) and indels. Variants used in downstream analyses were subjected to the quality control filter described in Wybouw, Kosterlitz, et al. (2019).

### 2.8 Principal component analysis

A principal component analysis (PCA) using the genomic data of the parental lines was conducted using the R package ‘SNPRelate’ (version 1.30.17) as described by Zheng et al. (2012). SNPs were first pruned (“set.seed(1000)”; “snpgdsLDpruning” function with “slide.max.bp = 50000”, “ld.threshold = 0.2” and “autosome.only = FALSE”)) before performing the PCA of the parental lines (“snpgdsPCA” function with autosome.only = FALSE). For each of the two BSA experiments, a PCA was created using the VCF file (section 2.7) in R (package *prcomp;* version 2.3.0), as described in (Snoeck, Kurlovs, et al., 2019). To do so, a correlation matrix containing individual SNP frequencies was used as input. Only SNP’s that differentiated the two parental lines from each BSA, and that were present on every treatment of their respective experiment (abamectin-selected and unselected replicates) were selected for the PCA. Two-dimensional PCA plots for each BSA experiment were created using ggplot2 (version 3.3.3) (Valero-Mora, 2010) in R.

### 2.9 Bulked segregant analysis mapping

Previously developed BSA methods (Kurlovs et al., 2019) were used to map loci associated with abamectin resistance in our experimental evolution assay using the “RUN_BSA1.02.py” script available at (https://github.com/rmclarklab/BSA). Statistical significance of the resulting BSA peaks was assessed using the permutation approach outlined by Wybouw, Kosterlitz, et al. (2019), where differences in allele frequencies between paired selected and control replicates were calculated iteratively with 1000 permutations and with a false discovery rate (FDR) < 0.05.

### 2.10 Predicted effects of genetic variants in coding sequences

To assess the potential effect of a variant allele on loci under selection identified in the BSA genomic scans, coding effects of SNPs and small indels identified by the GATK analysis (section 2.7) were predicted using SnpEff 5.0c (Cingolani et al., 2012), with a *T. urticae* coding sequence database derived from the June 23, 2016 annotation, available from the Online Resource for Community Annotation of Eukaryotes (ORCAE) (Sterck et al., 2012). The SNPsift toolbox, provided within the SNPeff package, was used per BSA experiment to filter the SNPeff output for variant alleles present only in the resistant parental line, absent in the susceptible parental line and enriched in all selected populations of each BSA (i.e., allelic depth (AD) of variant allele > AD of reference allele in selected samples).

### 2.11 Molecular analysis of *TuGlucl2* and its association with abamectin selection

The glutamate-gate chloride channel (GluCl) consists of five subunits, encoded by 5 to 6 different genes in *T. urticae* (Dermauw et al., 2012). In this section, we focussed on the *T. urticae GluCl2* gene (*TuGluCl2*) for downstream analyses. RNA was extracted from each of the parental lines as described in section 2.3. One μg of cDNA from each parental line was synthesised using the Maxima first strand cDNA synthesis kit (Thermo Fisher Scientific, Waltham, MA, USA). Primers Tu_GluCl2_dia_F (5’-TCATCGTCTCTTGGGTCTCC) and Tu_GluCl2_dia_R (5’-CCCATCGTCGTTGATACCTT), were used to amplify the fourth exon of *TuGluCl2* via a PCR reaction as described previously (Dermauw et al., 2012), using the cDNA of the four parental inbred lines as templates. The cycling conditions were set at 94 °C for 2 min with 30 cycles as following: 94 °C for 30 s, 54 °C for 45 s, 72 °C for 1 min and a final extension step of 5 min at 72 °C. A similar PCR-based amplification was also conducted using gDNA from the four parental lines, and from each treatment and control replicate of aBSA as templates. The PCR products were checked visually on 2% agarose gels, purified with the E.Z.N.A^©^ Cycle-Pure kit (Omega Bio-Tek, Norcross, GA, USA) and Sanger sequenced (LGC Genomics, Germany) with the PCR primers described above (‘Tu_GluCl2_dia’).

As the PCR amplification with the MAR-ABi template gDNA yielded multiple bands with the Tu_GluCl2_dia primers (section 3.5), we designed a new set of primers (‘TuGluCl2in’) based on the obtained partial sequence of the insert, targeting the largest amplicon: TuGluCl2in_F (5’-CGGGGCTTTACTTGAGTTTG) and TuGluCl2in_R (5’-CCCATCGTCGTTGATACCTT). We conducted a PCR reaction using the Expand Long Range PCR Kit (Roche, Basel, Switzerland) and the TuGluCl2in primers, with the gDNA of line MAR-ABi as template and the cycling conditions specified in the kit. Amplicons were visually inspected on a 2% agarose gel, purified with the E.Z.N.A^©^ Cycle-Pure kit (Omega Bio-Tek, Norcross, GA, USA) and Sanger sequenced with the TuGluCl2in primers, as well as with two internal primers (TuGluCl2internal_F: 5’-TAATTGGGCAAGACCTTGGA; TuGluCl2internal_R: 5’-TGGCAAAAGACAAAATCGAA). Open reading frame (ORF) finder (https://www.ncbi.nlm.nih.gov/orffinder/) was used to search this unexpected amplified sequence in *TuGluCl2* for potential protein encoding segments. In addition, we performed a Blastn search against the reference genome of the ‘London’ strain of *T. urticae* (Grbiċ et al., 2011) to investigate whether the sequence was present elsewhere in the genome.

To investigate whether the abamectin selected samples of aBSA were enriched with the *GluCl2* sequence present in the resistant parent MAR-ABi, DNA reads from all aBSA replicates, and parental lines MAR-ABi and SOL-BEi were mapped against a version of the three pseudochromosome *T. urticae* assembly, where we artificially replaced the London *GluCl2* gene sequence with the MAR-ABi *GluCl2* gene sequence. The mapping was performed as described in section 2.6. The resulting position-sorted bam files were used as input for GATK’s tool CollectReadCounts (version 4.1.7.0) to count DNA reads mapping specifically to the artificially introduced region in the genome for each of the samples.

## 3 RESULTS

### 3.1 Abamectin resistance in parental inbred lines and in F1 hybrid populations

Susceptibility to abamectin differed largely between parental lines (Table 1). MAR-ABi and ROS-ITi were 1200- and 200-fold more resistant to abamectin than SOL-BEi and JP-RRi, respectively. The susceptibility of F1 offspring relative to their parents showed that the inheritance of resistance is incompletely recessive in the aBSA mapping population, and incompletely dominant in the gBSA mapping population (Table 1).

**TABLE 1.** Toxicity bioassays of abamectin using the inbred parental lines of MAR-ABi, ROS-ITi, SOL-BEi, JP-RRi and F1 hybrid populations from aBSA and gBSA

|  | **Slope(±SE)** | **LC<sub>50</sub>(95% CI) / mg a.i. /L** | **χ<sup>2</sup>(df\*)** | **DD<sup>†</sup>** |
| --- | --- | --- | --- | --- |
| **aBSA** |  |  |  |  |
| SOL-BEi | 4.07 (±0.47) | 0.10 (0.08-0.12) | 34.37(18) |  |
| MAR-ABi | 0.96 (±0.20) | 118.8 (69.33-407.4) | 37.69(25) |  |
| F1: ♀ SOL-BEi x ♂ MAR-ABi | 1.52 (±0.16) | 2.27 (1.01-4.46) | 22.58(10) | -0.117 |
| **gBSA** |  |  |  |  |
| JP-RRi | 4.67 (±0.52) | 0.13 (0.12-0.14) | 4.96(18) |  |
| ROS-ITi | 4.31 (±0.60) | 30.59 (24.69-35.06) | 40.26(27) |  |
| F1: ♀ JP-RRi x ♂ ROS-ITi | 5.94 (±0.65) | 5.51 (5.02-6.00) | 21.86(17) | 0.373 |

\*df: degrees of freedom
†DD: degree of dominance. DD = 1: dominant; 1 > DD > 0: incompletely dominant; 0 > DD > -1: incompletely recessive; DD = -1: recessive).

### 3.2 Differential expression analysis between abamectin resistant and susceptible parental lines

A total of 1415 genes were differentially expressed (DEGs) (|Log_2_FC| > 2, padj < 0.05) in the abamectin resistant parent MAR-ABi versus the susceptible parent SOL-BEi of aBSA. Amongst these DEGs, 607 genes (42.9%) were upregulated and 808 genes were downregulated (57.1%) in MAR-ABi; 94 of the 1415 DEGs (6.64%) code for enzymes belonging to important detoxification gene families (CCEs, CYPs, DOGs, GSTs, SDRs and UGTs; Table S1). In gBSA, a total of 1268 genes were differentially expressed between the abamectin resistant parent ROS-ITi and the susceptible parent JP-RRi, of which 682 genes (53.8%) were upregulated and 586 genes (46.2%) downregulated; 70 of the 1268 DEGs (5.5%) code for important detoxification gene families (Table S2). Interestingly, the detoxification gene family of the Cytochrome P450s (CYPs) shows contrasting differential expression profiles between aBSA and gBSA. In the comparison of MAR-ABi versus SOL-BEi, the largest fraction (23 out of 29) of differentially expressed CYPs were upregulated (Table S1), whereas in the comparison of ROS-ITi versus JP-RRi the largest fraction (18 out of 24) of differentially expressed CYPs were downregulated (Table S2). Only CYP392E9 and CYP392A10v2 were upregulated in both experiments. Such contrasting gene expression profiles likely reflect the overall different life histories and acaricide selection regimes experienced by the parental lines (described in Kurlovs et al. (2022)), highlighting the need of an unbiased approach to associate genomic loci with abamectin resistance.

### 3.3 Experimental evolution of abamectin resistance

Crossing abamectin-susceptible and resistant lines successfully generated sufficient F1 female progeny to obtain a large mapping population for each BSA experiment. Subsequently, paired abamectin-selected and unselected control replicates were set-up as described in section 2.4. After the experimental evolution of these populations with and without abamectin selection, the survival of selected and control replicates was tested at a discriminating concentration of 15 mg a.i. /L for aBSA or 10 mg a.i. /L for gBSA. At these concentrations, the corrected survival of abamectin-selected populations was close to 100% (Figure 1), which was significantly higher than the corrected survival of control populations both in aBSA (F_1,69_ = 5614.8, p < 0.001) and in gBSA (F_1,77_ = 6094.4, p < 0.001). This indicates that the selected and unselected replicates differ largely in their susceptibility to abamectin and thus selection resulted in the evolution of abamectin resistance.

**FIGURE 1.**
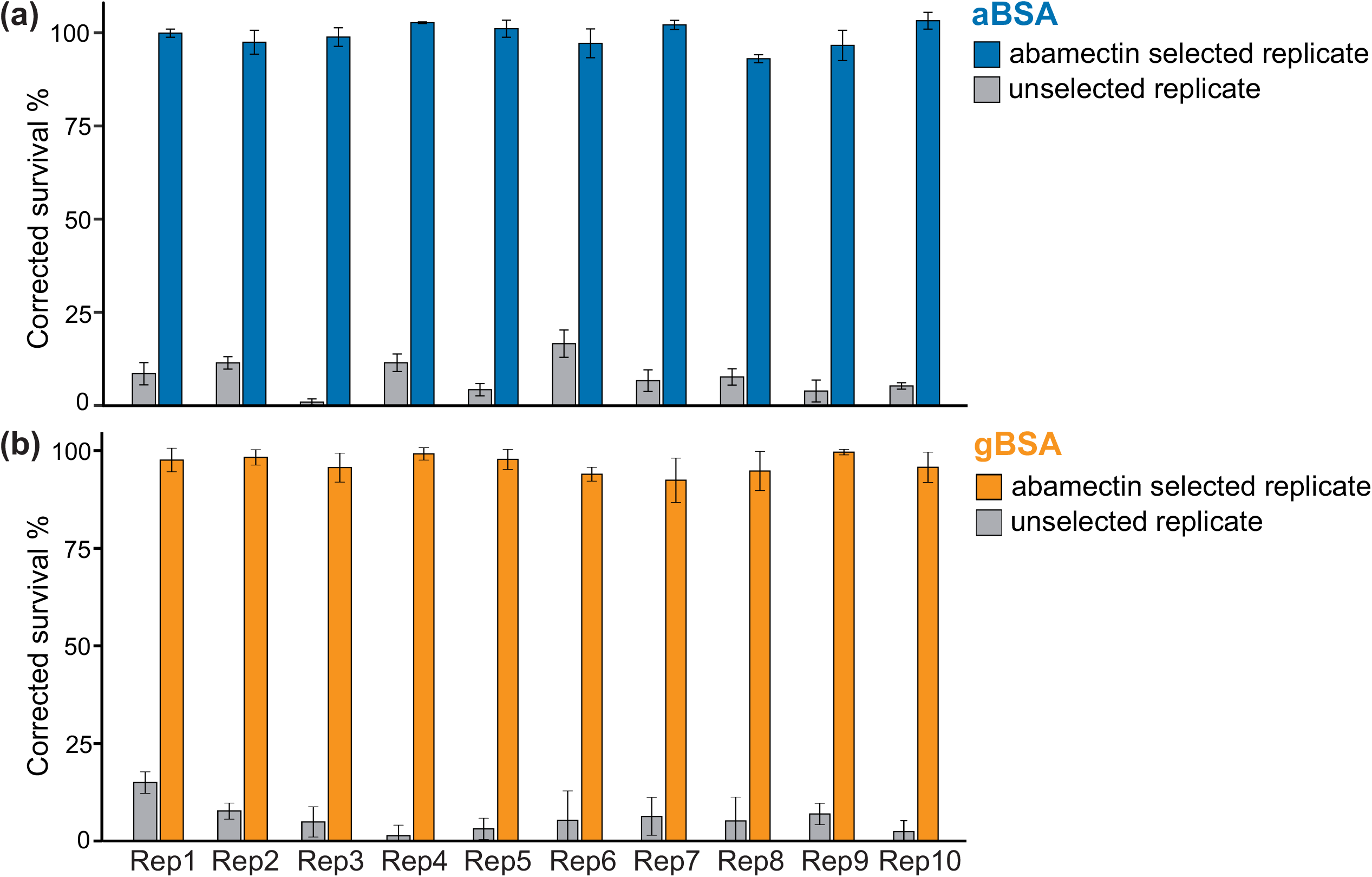
Susceptibility to abamectin of abamectin-selected and unselected control replicates from two independent BSA experiments, (a) aBSA and (b) gBSA. The corrected survival percentage of each replicate was scored in the adult stage after spraying with 15 mg a.i. /L and 10 mg a.i. /L abamectin for aBSA and gBSA, respectively. The survival of both sets of abamectin-selected replicates was higher than the survival of unselected replicates.

### 3.4 Genomic responses to abamectin selection and BSA analysis

After phenotyping the experimental populations in both BSA experiments, genomic DNA was extracted from each selected and unselected population and sequenced using the Illumina technology, resulting in an output of 29 to 45 million paired end reads of 100 bp long. For the parental strains, genomic DNA was extracted, sequenced and publicly released in the study of Kurlovs et al. (2022). Reads of all experimental populations for aBSA and gBSA and of the parental strains (MAR-ABi, ROS-ITi, SOL-BEi and JP-RRi) were aligned to the pseudochromosome genome assembly of *T. urticae* and subsequently used for variant calling (Wybouw, Kosterlitz, et al., 2019). For the parental strains, the PCA clearly separated the resistant parental lines from the susceptible parental lines along PC1, which explained 35.1% of the variation in the dataset (Figure 2a). PC2, explaining 32.7% of the variation, separated the susceptible parental lines used in the two BSA experiments (Figure 2a-b), while PC3 separated the two resistant parental lines (Figure 2b). For both experiments, aBSA and gBSA, the resulting variant call file (VCF) (available as Data S1_aBSA_vcf and S2_gBSA_vcf, respectively on figshare XXXX) were used to identify high quality SNPs discriminating MAR-ABi from SOL-BEi in aBSA, and ROS-ITi from JP-RRi in gBSA. For aBSA, a total of 645,199 high-quality, segregating SNPs were merged into a correlation matrix to analyse the global genomic responses to abamectin selection. The resulting PCA explained 71.5% of the variation in the dataset along PC1, and 2.8% along PC2 (Figure 2c). For gBSA, a total of 569,642 high-quality, segregating SNPs were merged into the correlation matrix. The resulting PCA plot explained 63.7% of the variation in the dataset along PC1, and 6.0% along PC2 (Figure 2d). In both PCA plots, the unselected controls and selected replicates clearly clustered into two distinct groups along PC1, while the replicates of gBSA were more heterogeneously spread along PC2 than the replicates of aBSA (Figure 2c-d).

**FIGURE 2.**
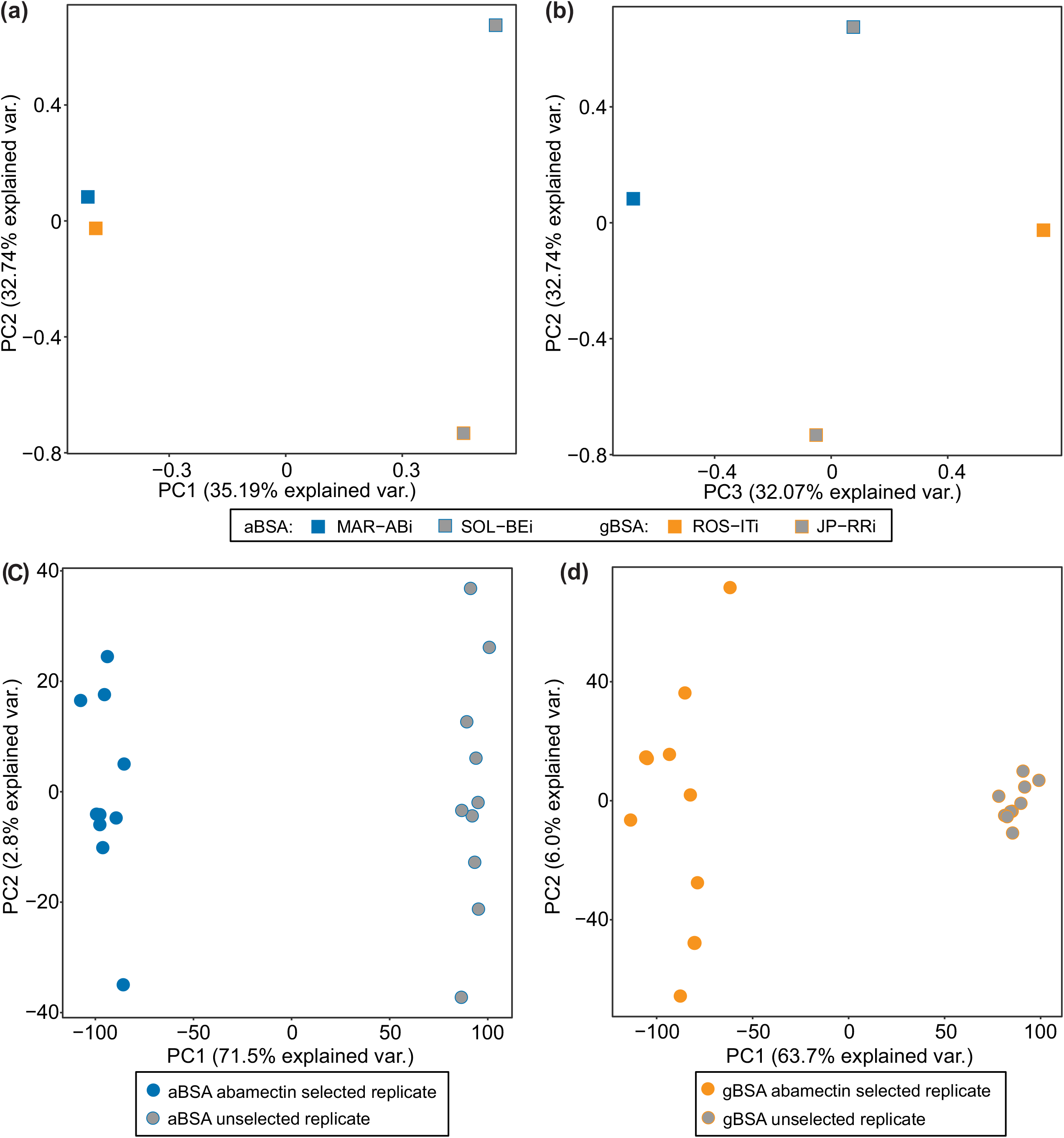
Selection for abamectin resistance is associated with a genomic response. a-b) Principal component analysis (PCA) of the parental inbred lines used in the BSA mapping experiments. PC1 evidently separated the resistant lines (MAR-ABi and ROS-ITi) from the susceptible lines (SOL-BEi and JP-RRi), while PC2 separated the two susceptible parental lines, and PC3 separated the two resistant parental lines. c) PCA with unselected and abamectin-selected replicates of aBSA based on genome-wide allele frequencies at polymorphic sites, where PC1 clearly separated unselected and selected replicates. d) PCA with unselected and abamectin-selected replicates of gBSA based on genome-wide allele frequencies at polymorphic sites, where PC1 clearly separated unselected and selected replicates. Individual replicates are coloured according to the treatment group (legend).

Significant deviations in allele frequency between abamectin-selected and unselected replicates were used to determine local regions that responded to selection along sliding windows across the genome (Kurlovs et al. 2019). Four QTL peaks associated with abamectin selection in aBSA, and three QTL peaks in gBSA, exceeded the genome-wide significance threshold (FDR < 0.05). Notably, the parental haplotypes nearly reached fixation at each of the four QTL peaks (Figure 3a). Allele frequencies of all QTL peaks reflected selection in the direction of the abamectin-resistant parent. As observed in the PCA plots (Figure 2), genome-wide variation between replicates of gBSA was higher than between replicates of aBSA (Figure S1). The three QTL peaks of gBSA overlapped with three of the four QTL peaks of aBSA, with an offset between peaks of the two BSA experiments ranging from 20kb to 440 kb (Figure 3). A 500kb region surrounding the top significant QTL window of each peak was further analysed. The differential expression analysis of the genes under this 500kb window showed that most of these candidate genes were not differently expressed between resistant and susceptible parental lines (MAR-ABi vs. SOL-BEi and ROS-ITi vs. JP-RRi), with |log2FC| ≥ 2 and p-adj < 0.05 (Table S3-S7). QTL peaks were named according to their position in aBSA (QTL1-4). Thus, aBSA contained QTL peaks 1-4, while gBSA contained QTL peaks 2-4.

**FIGURE 3.**
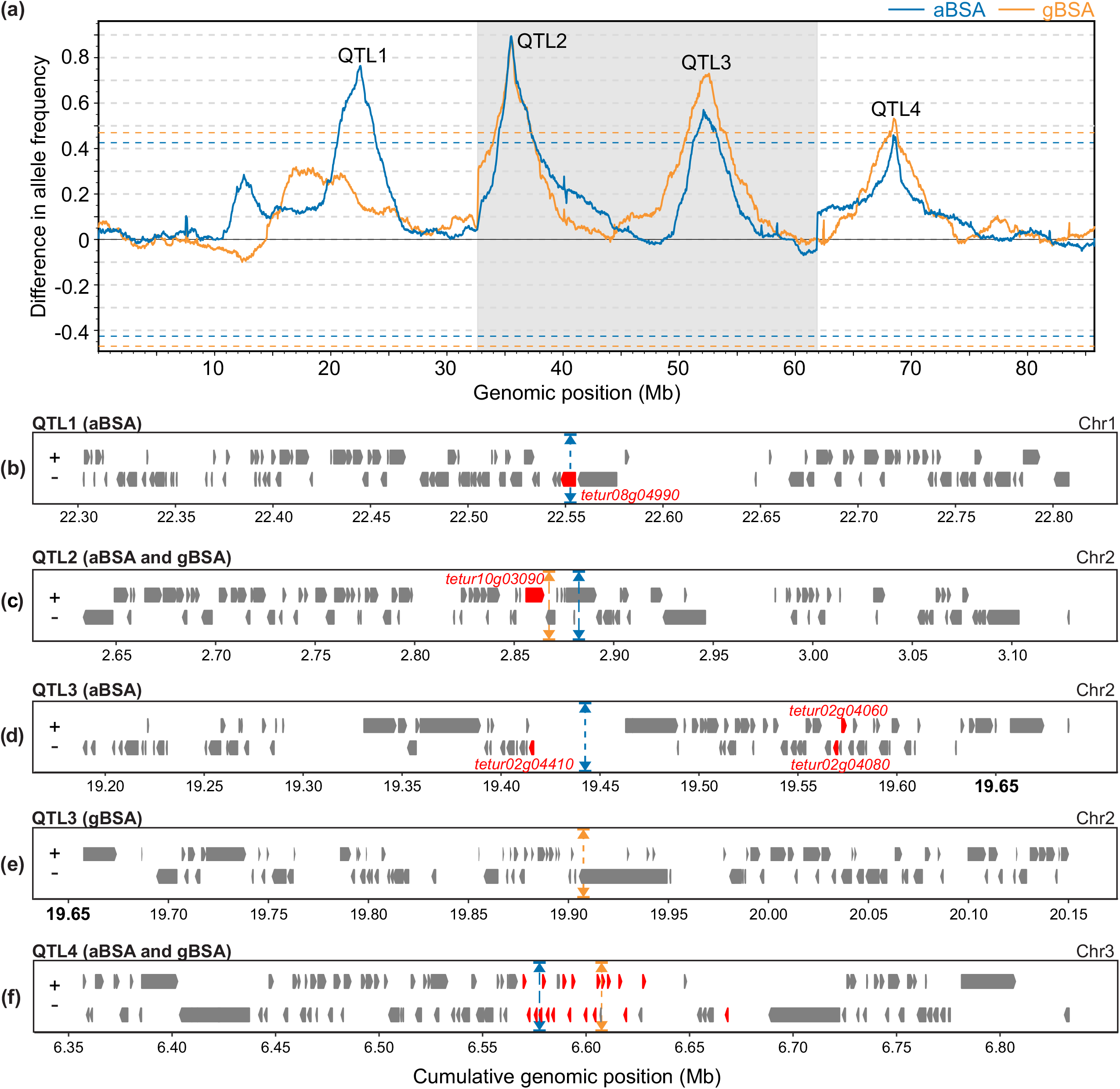
Genomic responses to abamectin selection. (a) Averaged genome-wide allele frequency differences between abamectin-selected and unselected replicates in aBSA (blue line) and gBSA (orange line). QTL associated with abamectin resistance (QTL 1-4) show as peaks that deviate from the genome-wide average. Dashed lines delineate the statistical threshold for QTL detection (FDR < 0.05). (a-f) Genomic positions are denoted on the x-axis, with the three-pseudochromosome (Chr1-3) configuration of *T. urticae* indicated by alternating shading. (b-d) Candidate genes within 500 kb genomic brackets around QTL peaks. Triangles positioned along the top and bottom boundaries of each plot represent the top genomic window, averaged for each BSA experiment: aBSA (blue), gBSA (orange). The orientation of the gene models is indicated by “+” or “-” for forward and reverse strands, respectively. Putative candidate genes within the QTL peaks are highlighted in red. (b) QTL peak 1: glutamate-gated chloride channel 2 (*TuGluCl2: tetur08g04990);* (c) QTL peak 2: glutamate-gated chloride channel 3 (*TuGluCl3: tetur10g03090);* (d-e) QTL peak 3: glutamate-gated chloride channel 1 (*TuGluCl1*: *tetur02g04080*) and DEAD/DEAH box DNA helicases (*tetur02g04060, tetur02g04410);* (f) QTL peak 4: multiple chemosensory receptors (Table S7).

QTL peak 1 was located at 22.55 Mb on pseudochromosome 1, and only occurred in aBSA (Figure 3a, b). One of the genes forming the target-site of abamectin, the glutamate-gated chloride channel subunit 2 (*TuGluCl2 [tetur08g04990]*), was located at the centre of the peak (Figure 3b, Table S3). We did not find any allele variants that could impact the function of *TuGluCl2* in any of the abamectin-selected replicates.

QTL peak 2 was located approximately at ~2.9Mb on pseudochromosome 2, with the top peaks from each BSA experiment only 20kb away from each other (Figure 3a). An allele previously confirmed to confer target-site insensitivity to abamectin, G326E in *GluCl3* (*tetur10g03090*), was located approximately within 40kb of each peak (Figure 3c, Table S4) (Dermauw et al., 2012; Mermans et al., 2017; Xue et al., 2020). The frequency of the G326E mutation was higher in the selected replicates than in the unselected replicates of both BSA experiments (Table S8). In addition, a V273I substitution belonging to the parent MAR-ABi was found in the transmembrane domain 1 (TM1) of *GluCl3*, and the frequency of this mutation was higher in the selected replicates than in the unselected replicates of aBSA (Figure 4a, Table S8).

**FIGURE 4.**
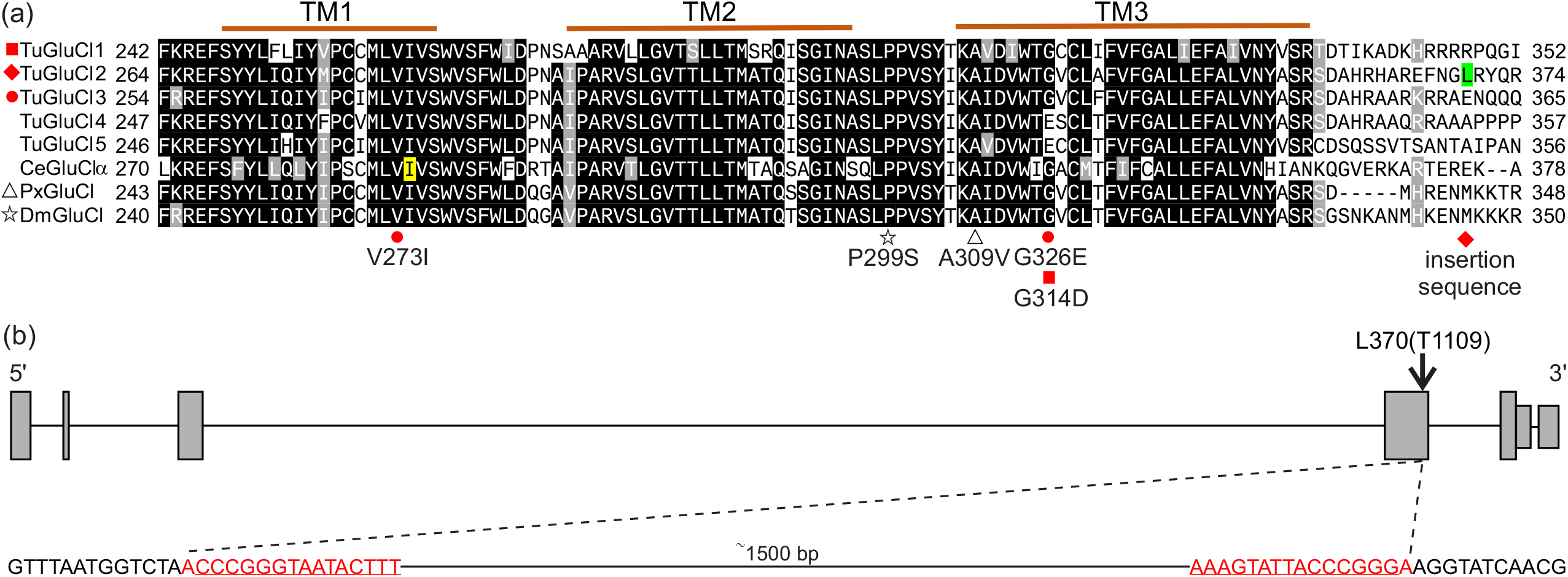
Sequence and gene model of the glutamate-gated chloride channel. (a) Alignment of transmembrane domains 1, 2 and 3 (TM1, TM2 and TM3) of GluCl genes in *T. urticae* (Tu), *C. elegans* (Ce) *P. xylostella* (Px) and *D. melanogaster* (Dm). An 80% threshold was used for identity (black background) or similarity shading (grey background). The delineation of TMs is based on the crystal structure of the *C. elegans GluClα*. A predicted ivermectin binding site at position I290 from a three-dimensional structure of *GluClα* in *C. elegans* is shaded in yellow. The red boxes and dots indicate the point mutations found in this study. The position of the insertion sequence in *GluCl2* is shaded in green and indicated with a red rhombus. The P299S substitution in TM2-TM3 and the A309V in TM3 that have been previously associated with avermectin resistance in *D. melanogaster* and *P. xylostella* are indicated by stars and triangles, respectively. (b) The insertion sequence in *GluCl2* of the resistant line MAR-ABi. The dashed line indicates the location of the insertion after TM3 in *GluCl2*. Grey large rectangles, small rectangles and lines respectively represent exons, untranslated regions (UTR) and introns. The underlined bases represent a terminal inverted repeat pair.

QTL peak 3 was located between 19.45Mb and 19.92Mb on pseudochromosome 2 of aBSA and gBSA, respectively. Another subunit of the glutamate-gated chloride channel, *TuGluCl1*, was located within QTL peak 3 in aBSA (~100kb from the top peak), but it was located more distantly in gBSA (~330kb from the top peak; Figure 3a). A G314D variant in *TuGluCl1* was identified in both experiments (Figure 3d-e, Figure 4a, Tables S5-S6). The frequency of this mutation was higher in the selected replicates than in the unselected replicates in both experiments (Table S8). In addition, two genes, *tetur02g04410* and *tetur02g04060*, both annotated as a DEAD/DEAH box DNA helicase were located 40 and 130 kb away from QTL peak 3 of aBSA, respectively (Figure 3d, Table S5).

QTL peak 4 was located at ~6.6Mb on pseudochromosome 3, with the top peaks from each BSA 50kb away from each other (Figure 3a). Among the 108 annotated genes found to overlap in the 470kb collective regions of QTL peak 4 of aBSA and gBSA, two genes encoding degenerin/Epithelial Na+□ channels (ENaCs) and 19 genes encoding chemosensory receptors were found (Figure 3f, Table S7). In addition, at the edge of the range around QTL peak 4, two functional CYP genes and two functional CCE genes were found (Figure 3e). The CCE genes (*TuCCE15* and *TuCCE50*) showed significant differences in expression levels when comparing MAR-ABi versus SOL-BEi, with *TuCCE15* being downregulated (Log_2_FC = −2.38) and *TuCCE50* being moderately upregulated (Log_2_FC = 1.39; Table S7).

### 3.5 *GluCl2* genotyping

Given that *GluCl2* was found at the centre of QTL peak1 of aBSA (Figure 3a-b, Table S3), but no allele variants could be identified that differed between the abamectin-selected and the unselected control replicates, we further investigated the genome of the resistant parent MAR-ABi. A PCR with a previously published primer pair (Dermauw et al. 2012) and MAR-ABi gDNA template resulted in multiple amplicons for *GluCl2* (Figure S2a). Further, a long-range PCR with newly designed primers revealed a 1547 bp long insertion located between transmembrane domain 3 (TM3) and TM4 within the fourth exon of *GluCl2* (Figure 5, Figure S2c, Supplementary file 1). The fourth exon of *GluCl2* is expressed in all parental lines except in MAR-Abi, as shown by PCR amplification of cDNA of the parental strains (Figure S2b). This was further confirmed by a *de novo* assembled transcriptome, which shows that the insertion causes a premature stop on the fourth exon of *GluCl2* in MAR-ABi (Figure 4, Figure S3).

The insertion was present in all abamectin-selected populations of aBSA (Figure S2d) and nearly reached fixation based on the frequency of the MAR-ABi allele at QTL peak 1 (Figure S1a). In addition, the number of DNA read counts mapping to the *GluCl2* insertion sequence was considerably higher in the abamectin-selected replicates of aBSA than in the control replicates, and much higher in MAR-ABi than in SOL-BEi (Figure S4). The insertion contains short terminal inverted repeats (TIRs) of 15bp, and it either lacks the usual target site duplications associated with transposable elements, or these duplications are very short (i.e., 2bp flanking the insertion; Figure 4b). We did not find any open reading frames within the insertion, and accordingly, it does not encode any proteins. It also does not match previously described transposon families according to queries using BLASTn, BLASTx or Dfam databases. The insertion was not found anywhere else in the reference genome of *T. urticae*.

## 4 DISCUSSION

While a plethora of mechanisms underlying abamectin resistance have been identified in arthropod and non-arthropod taxa, high abamectin resistance levels could, in most cases, not be attributed to one single mechanism (Choi et al., 2017; Dermauw et al., 2012; Ghosh et al., 2012; Khan et al., 2020; Kwon et al., 2010; Mermans et al., 2017; Riga et al., 2014; Xue et al., 2020, 2021). Mutations in genes encoding subunits of the glutamate-gated chloride channel (*TuGluCl1* and *TuGluCl3*) are major factors contributing to abamectin resistance in *T. urticae* (Dermauw et al., 2012; Kwon et al., 2010; Mermans et al., 2017; Xue et al., 2020). However, introgression experiments of single target-site mutations have revealed that they do not confer a strong resistance phenotype without additive or synergistic effects of other resistance factors. In addition, the presence of target-site mutations can result in pleiotropic effects that negatively impact the fitness of spider mites without abamectin application (Bajda et al., 2018). Together, this suggests that a complex and polygenic architecture underlies the often stable phenotype of resistance found in laboratory and field populations (Xue et al., 2020). Here, we investigated the genetic basis of abamectin resistance in two genetically unrelated populations of the two-spotted spider mite *T. urticae*. Corroborating a complex and polygenic architecture, we found multiple loci associated with abamectin resistance (Figure 1–3). In addition, investigating two lines derived from genetically unrelated, resistant *T. urticae* populations provided important insights into the intraspecific diversity of mechanisms associated with the evolution of this phenotype.

Different modes of inheritance of abamectin resistance have been found across populations of *T. urticae*, including incompletely recessive, recessive, or incompletely dominant (Dermauw et al., 2012; He et al., 2009; Kwon et al., 2010; Salman & Ay, 2009). Here, independent crosses of two resistant lines with a susceptible line each yielded low levels of resistance to abamectin in the F1 generation, one and two orders of magnitude lower than the resistant parents of gBSA and aBSA, respectively (Table 1). The level of resistance of the segregating mapping population of gBSA was almost twice as high than the aBSA population, despite resistance in MAR-ABi being approximately four times higher than resistance in ROS-ITi. These differences in the level of resistance in the F1 generation indicate an incompletely recessive or incompletely dominant mode of inheritance (Table 1). Furthermore, the difference in the magnitude of resistance between the mapping populations of each BSA may also be indicative of differences in the mechanisms underlying resistance in the two parental lines.

We conducted two independent BSA experiments to identify the loci associated with abamectin resistance without any prior hypothesis. For each experiment, we performed genome-wide scans using ten abamectin-selected and ten unselected populations to calculate allele frequency differences along overlapping genomic windows (Figure 3, Figure S1). The scans revealed a polygenic basis associated with abamectin resistance, and despite using genetically unrelated mite lines in this study (Figure 2a-b), we found a striking overlap of the QTL peaks between BSA experiments (Figure 3a). Abamectin selection resulted in three QTL peaks in gBSA, which overlapped with three of the four QTL peaks found in aBSA. Variation was larger between the selected replicates of gBSA than in any other treatment or control group, as evidenced in the PCA plot (Figure 2c-d). Large variation between the selected replicates of gBSA also resulted in more coarsely defined QTL peaks than in aBSA, because each of the replicates of gBSA was further located from the average, when compared to aBSA (Figure S1). The larger variation among replicates in gBSA, and thus the coarser mapping resolution in the genomic scans, might be the result of stronger population bottlenecks experienced by gBSA replicates, which were possibly caused by a stronger selection regime than in aBSA. Population bottlenecks can lead to fewer sampled recombination events due to low population sizes, and thus increase the variation among replicates. The larger variation in gBSA compared to aBSA also meant that the average of each QTL peak differed by a few kb between BSA experiments (Figure 3a). This mismatch was largest in QTL peak 3, which differed by approximately 500kb between experiments (Figure 3d-e). Furthermore, one of the QTL peaks found in aBSA was not present in gBSA, QTL peak 1, indicating a different genetic basis to abamectin resistance in each of the parental lines. Nonetheless, the three shared QTL peaks found independently suggest that the evolution of abamectin resistance involves a polygenic and likely complex genetic basis, in which multiple genes either work additively or synergistically to obtain high levels of resistance.

Within QTL peaks 2 and 3 of aBSA resided two genes that encoded subunits of the GluCl channel, *TuGluCl3* and *TuGluCl1*, respectively (Tables S4, S5). Previous work demonstrated the role of the G326E and G314D mutations (in *TuGluCl3* and *TuGluCl1* respectively) in conferring resistance to abamectin (Dermauw et al., 2012; Kwon et al., 2010). *TuGluCl3* was found within QTL peak 2 in both BSA experiments, and in all abamectin-selected replicates the reported G326E target-site mutation was enriched (Figure 3c, Table S4, Table S8). In addition, we found a V273I mutation in *TuGluCl3* present in the MAR-ABi parent and enriched in the abamectin-selected replicates of aBSA (Figure 4a, Table S5, Table S8). This mutation is adjacent to a predicted ivermectin binding site at position I290 from a three-dimensional structure of *GluClα* in *C. elegans* (Ludmerer et al., 2002). The G314D mutation in *TuGluCl1*, labelled as G323D by Kwon et al. (2010), was found in higher frequencies in all the abamectin-selected replicates of aBSA and gBSA than in unselected controls (Table S8). *TuGluCl1* was found ~100kb and ~300kb away from the top of QTL peak 3 in aBSA and gBSA, respectively (Figure 3a, d-e). It is possible that lower variation among replicates in gBSA would have resulted in this mutation appearing closer to the QTL peak. However, *TuGluCl1* was also located far away from the QTL peak 3 in aBSA in comparison to similar BSA studies, where causal genes have been found within tens of kb of the QTL peak (Fotoukkiaii et al., 2021; Snoeck, Kurlovs, et al., 2019; Wybouw, Kosterlitz, et al., 2019). Hence, we hypothesized that other genes than those encoding the target-site might play a role in abamectin resistance, including two genes at QTL peak 3 annotated as a DEAD/DEAH box DNA helicase (Table S3). Among the six major super families of described helicases, members of the DEAD/DEAH box complex are part of superfamily 2, which are generally involved in the unwinding of nucleic acids and in the metabolism of RNA molecules. DEAD/DEAH box helicases in RNA viruses are essential for synthesis of new genomic RNA (Gilman et al., 2017). Recently, ivermectin, which is structurally almost identical to abamectin (Lespine, 2013), has been identified to interfere with the replication process of flavoviruses and coronaviruses by targeting DEAD-box helicase activity (Caly et al., 2020; Mastrangelo et al., 2012). Interestingly, both *T. urticae* DEAD/DEAH box helicase genes near QTL peak 3 were downregulated in the resistant line MAR-ABi (Table S3). DEAD/DEAH box helicase genes were also shown to be downregulated in the nematode *Brugia malayi* after exposure to ivermectin (Ballesteros et al., 2016). Whether mutations in the sequence of the DEAD/DEAH box DNA helicases found in this study alter their expression or function, and how this might be linked to abamectin resistance, remains to be investigated.

Another GluCl subunit, *TuGluCl2*, was mapped under QTL peak 1 in aBSA, but not in gBSA (Figure 4a, Table S1). Previously, mutations associated with target-site resistance were not found in *TuGluCl2* (Dermauw et al., 2012). However, we found that the parent MAR-ABi, as well as all the abamectin-selected replicates, contained an insertion of ~1500bp in the fourth exon of *TuGluCl2*, between transmembrane domain 3 and 4 (Figure 4, Figure S3, Supplementary file 1). The characteristic short terminal inverted repeats suggest that the insertion could be a non-autonomous miniature inverted transposable element (MITE)(Lu et al., 2012; Wicker et al., 2007), but as it is larger than other MITEs reported and it seems to lack target site duplications, its precise classification warrants further investigation. RNA expression levels showed, in contrast to SOL-BEi, that *GluCl2* is not expressed in the parental line MAR-ABi (Figure S2b). This suggests that *GluCl2* is likely not functional in MAR-ABi, and that abamectin application selects for the non-functional variant of *GluCl2*.

Since both G326E and G314D mutations in *TuGluCl1* and *TuGluCl3* were selected in all replicates of aBSA, along with the insertion in *TuGluCl2*, we can suspect that the disruption of a subunit of *GluCl2* has an adaptive value to the MAR-ABi line. This is an interesting observation, as the joint action of the G326E and G314D mutations in *TuGluCl3* and *TuGluCl1* respectively have been shown to be confer higher resistance to abamectin than each mutation alone in *T. urticae* (Riga et al., 2017), but the additional presence of the disrupted *TuGluCl2* variant could potentially explain the extremely high levels of resistance to abamectin of the line MAR-ABi compared to ROS-ITi (Table 1). Reducing the expression or completely knocking-down a subunit that is susceptible to the pesticide might reduce overall susceptibility to abamectin, potentially by forming an heteromeric ion channel only with those GluCl subunits that harbour resistance mutations (see Figure 1 from Xue et al., 2021). Similarly, nematodes have also evolved multiple copies of the GluCl gene (O’Halloran, 2022) and *C. elegans* native GluCls are believed to be composed of two to five different subunits (Wolstenholme & Neveu, 2022). In *C. elegans*, simultaneous mutation of three genes encoding glutamate-gated chloride channel a-type subunits has been shown to confer high-level resistance ivermectin, while mutating any two channel genes confers only modest or no resistance (Dent et al., 2000). In addition, it was reported that a naturally-occurring amino-acid deletion in the alpha subunit of in *C. elegans GluCl channel 1* results either in the decreased expression of the gene or in alterations to the chromatin structure around its sequence, conferring resistance to avermectin compounds (Evansid et al., 2021; Ghosh et al., 2012; Jones & Sattelle, 2008). Gene knock-out, although rarely reported, has been previously documented and experimentally linked to spinosad and neonicotinoid resistance in *Drosophila* and in *Plutella xylostella*, acting via the disruption of different nicotinic acetylcholine receptor subunits (reviewed in Feyereisen et al., 2015). Non-sense mutations and transposable elements disrupting gene function associated with resistance are only possible when a degree of functional redundancy is present (Baumann et al., 2010; Wilson, 1993; Wilson & Ashok, 1998). Whether the disruption of GluCl subunits has, for example, an impact on the fitness costs associated with harbouring the G326E (*GluCl3*) and G314D (*GluCl1*) mutations (Bajda et al., 2018) is a possibility that remains to be explored empirically. In this context, it is interesting to remark that inbreeding fixed the inserted sequence in *TuGluCL2* of the line MAR-ABi, which was only present at low frequency in the parental MAR-AB line, as this strain showed the 281bp diagnostic band for *TuGluCl2* (Dermauw et al. 2012), even when this line was under frequent abamectin selection, which could indicate a fitness cost.

Within QTL peak 4, multiple chemosensory receptors (CRs) were found (Table S7). Chemosensory receptors are major determinants of host plant acceptance in arthropods, and more than 400 intact gustatory receptors have been identified in *T. urticae* (Ngoc et al., 2016; Wicher & Marion-Poll, 2018; Wybouw, Kosterlitz, et al., 2019). As such, their potential as novel targets for pest control in crops is rising (Venthur & Zhou, 2018). Given their position in the QTL, it is possible that CRs are associated with abamectin resistance in *T. urticae* (Table S7). If so, binding of abamectin to CRs might lead to the activation of detoxification pathways, and they might hence act as xenosensors (Ingham et al., 2020). In nematodes, the transcription factor *cky-1* has been linked with ivermectin resistance, as this gene is within a locus under selection in ivermectin-resistant populations worldwide, and functional validation using knockdown experiments support the observation that *cky-1* is associated with ivermectin survival (Doyle et al., 2022). In mites, transcriptional regulation could lead to the modular control and differential expression of groups of genes, and at least be partially responsible for some of the genes that are differentially expressed between parental lines. It was recently shown that especially P450s and DOGs are *trans*-regulated in some of these lines (Kurlovs et al., 2022). In addition, it has been shown that the P450 *Cyp392A16* from *T. urticae* could metabolize abamectin to a non-toxic metabolite (Papapostolou et al., 2022; Riga et al., 2014), further suggesting that increased expression via a yet to be identified regulator in QTL4 could contribute to resistance.

## 5 CONCLUSION

High-resolution QTL mapping revealed the polygenic basis of abamectin resistance in two unrelated populations of the two-spotted spider mite *T. urticae*. We found three similar loci, and also one additional locus associated with resistance. Previously documented resistance mutations in genes encoding subunits of the glutamate-gated chloride channel (GluCl), the target-site of abamectin, were mapped in two independent BSA experiments. We also found that abamectin selects for a variant of GluCl subunit 2 that is very likely not functional, thus providing one of the rare examples were gene disruption of a target-site confers resistance. In addition, novel candidate loci associated with abamectin resistance were found, such as DNA helicases and chemosensory receptors. Our parallel experimental evolution set-up unravelled differences in the genetic mechanisms underlying the resistance phenotype between genetically distinct populations of a cosmopolitan species, hence, efforts to identify resistance mutations in field populations need to account for the diversity of resistance mechanisms in populations. Furthermore, our study opens the possibility to investigate how these variants are maintained in the field, and the different evolutionary origins and consequences of intraspecific variants underlying resistance.

## 6 ACKNOWLEDGEMENTS

WXX is the recipient of a doctoral grant from China Scholarship Council (CSC). This work was supported by the European Union’s Horizon 2020 research and innovation program [ERC consolidator grant 772026-POLYADAPT, and 773902-SuperPests].

## 7 DATA AVAILABILITY

All the sequence data generated in this study, including the parental lines and the segregating populations have been submitted to the Sequence Read Archive (SRA) and accession numbers will be provided upon publication.

## 8 CONFLICT OF INTEREST

The authors declare no conflict of interest related to this manuscript.

## 12 SUPPLEMENTARY TABLES

**Table S1.** Differentially expressed genes (Log2 Fold Change (Log2FC) > |2|, padj <0.05) in the comparison of the resistant parent MAR-ABi and the susceptible parent SOL-BEi of experiment aBSA

**Table S2.** Differentially expressed genes (Log2 Fold Change (Log2FC) > |2|, padj <0.05) in the comparison of the resistant parent ROS-ITi and the susceptible parent JP-RRi of experiment gBSA

**Table S3.** Genes within a 500kb bracket around the top genomic window of QTL peak 1 in aBSA, located at ~22.5Mb on Chr1

**Table S4.** Overlapping genes within a 500kb bracket around the top genomic window of QTL peak 2 in aBSA and gBSA, located at ~2.8Mb on Chr2

**Table S5.** Genes within a 500kb bracket around the top genomic window of QTL peak 3 in aBSA, located at ~19.4Mb on Chr2

**Table S6.** Genes within a 500kb bracket around the top genomic window of QTL peak 3 in gBSA, located at ~19.9Mb on Chr2

**Table S7.** Overlapping genes within a 500kb bracket around the top genomic window of QTL peak 4 in aBSA and gBSA, located at ~6.6Mb on Chr3

**Table S8:** Allele frequency of resistance mutations in *TuGluCl1* and *TuGluCl3* in all replicates and parental strains of aBSA and gBSA

## 13 SUPPLEMENTARY FIGURES

**FIGURE S1.** Genome-wide allele frequencies (n=10) of the resistant parent in abamectin-selected replicates of aBSA (blue lines) and gBSA (orange lines), and unselected control replicates (grey lines).

**FIGURE S2.** 2% agarose gel electrophoresis with PCR amplicons of the fourth exon of *GluCl2*. (a) PCR screening with gDNA from the four parental inbred lines shows a diagnostic 281 bp band for *GluCl2*, and a >1500 bp band when an insertion is present, as in line MAR-ABi. (b) PCR screening with cDNA from the three parental lines SOL-BE, ROS-ITi and JP-RRi shows the diagnostic 281 bp band of *GluCl2*, which is absent in line MAR-ABi. (c) PCR screening of the full-length insertion sequence using gDNA from the MAR-ABi line. (d) PCR screening with gDNA from each abamectin-selected (ABA; top panel) and unselected control replicates (CON; bottom panel) from aBSA. L: 100bp DNA ladder (Promega).

**FIGURE S3.** *GluCl2* gene from several *Tetranychus urticae* lines. Alignments of a portion of *GluCl2* using (a) nucleotide sequences and (b) amino acid sequences from abamectin - susceptible and -resistant parental lines (SOL-BEi and MAR-ABi) and the reference strain London. Red arrows indicate the start of the insertion in line MAR-ABi, which causes a premature stop. Sequences shared between the three lines have a black background (>90% similarity), while sequences shared between two lines have a grey background.

**FIGURE S4.** Number of DNA reads from the genomes of the parental lines (MAR-ABi and SOL-BEi) and the experimental populations of aBSA (abamectin-selected and unselected replicates) that mapped to the insertion sequence in *GluCl2*.

## Notes

### Competing Interest Statement

The authors have declared no competing interest.

