## Supplementary figures and images for "Intraspecific diversity in the mechanisms underlying abamectin resistance in a cosmopolitan pest"

### Figure S1

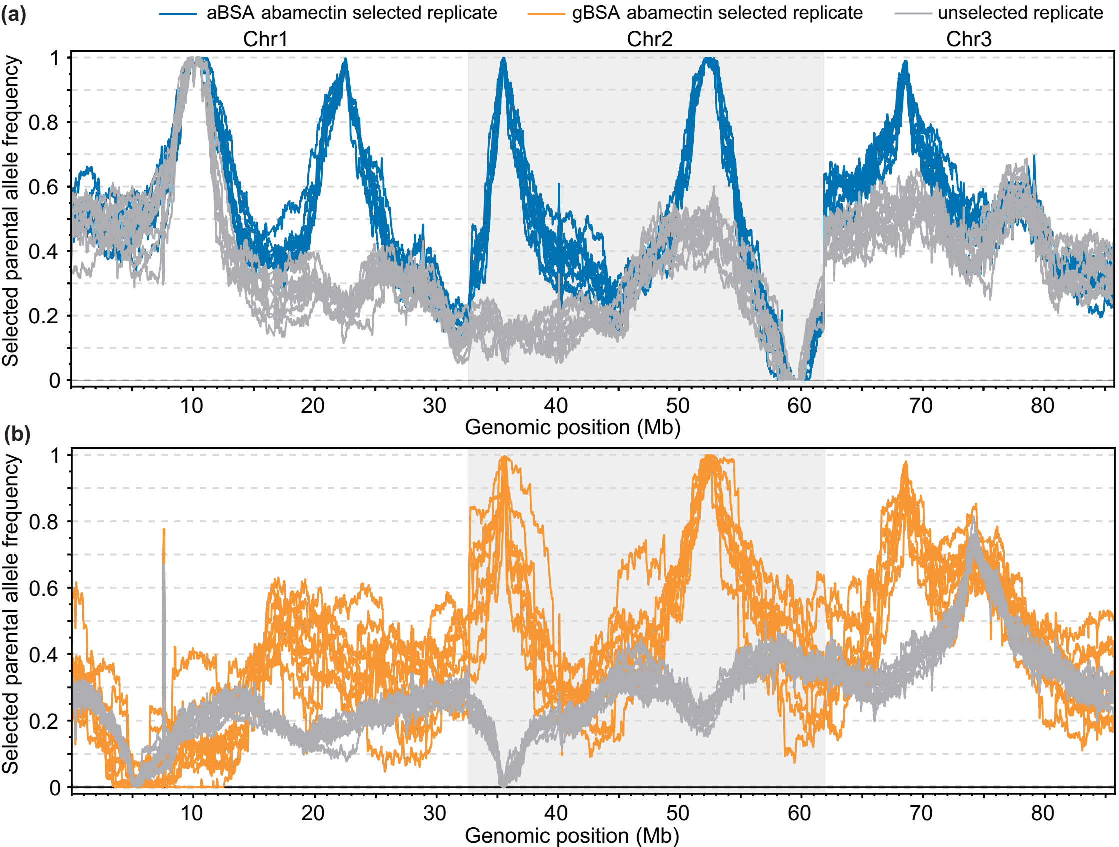

### Figure S2

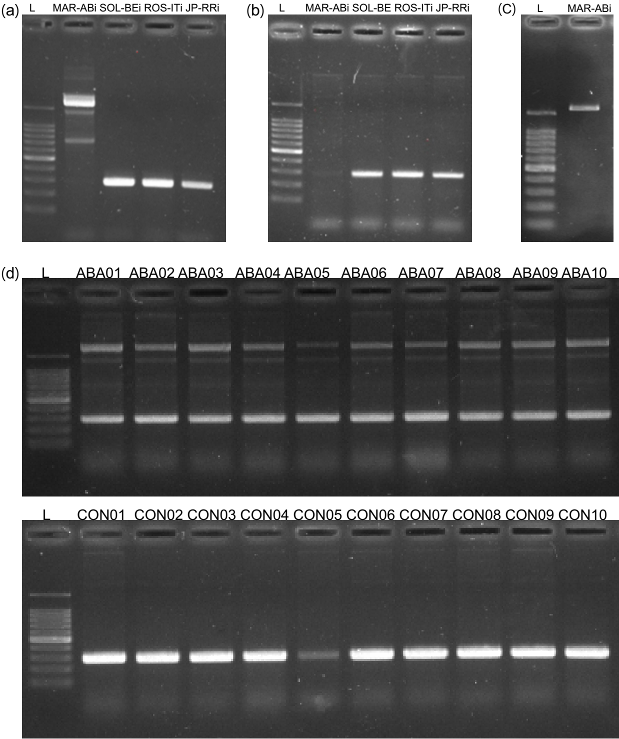

### Figure S4

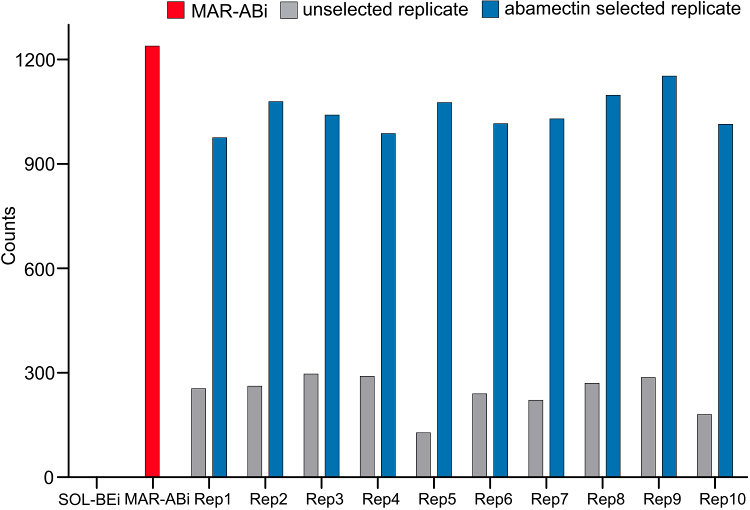
