## Supplementary material for "Intraspecific diversity in the mechanisms underlying abamectin resistance in a cosmopolitan pest": Figure S3

**(a)**

```
London  GluCl2 TTTGTTTTCGGGGCTTTACTTGAGTTTGCCTTGGTAAACTATGCTTCACGAAGTGATGCT 1080
SOL-BEi GluCl2 TTTGTTTTCGGGGCTTTACTTGAGTTTGCCTTGGTAAACTATGCTTCACGAAGTGATGCT 1080
MAR-ABi GluCl2 TTTGTTTTCGGGGCTTTACTTGAGTTTGCCTTGGTAAACTATGCTTCACGAAGTGATGCT 1080

London  GluCl2 CATCGCCATGCCAGAGAGTTTAATGGTCTAAGGTATCAACGACGATGG-GATAGGGATGG 1139
SOL-BEi GluCl2 CATCGCCATGCCAGAGAGTTTAATGGTCTAAGGTATCAACGACGATGG-GATAGGGATGG 1139
MAR-ABi GluCl2 CATCGCCATGCCAGAGAGTTTAATGGTCTAACCCGGGTAATACTTTTCAGCATTTCATGG 1140

                          inserted site in MAR-ABi

London  GluCl2 TAATGTTATCTCGGATGAAACCTCTTATGCACTGAGGCCATTGGTTATCAAAGGAAGTG- 1198
SOL-BEi GluCl2 TAATGTTATCTCGGATGAAACCTCTTATGCACTGAGGCCATTGGTTATCAAAGGAAGTG- 1198
MAR-ABi GluCl2 CAA-----GCTATGACCAAACGAG-CAAGCCTTGCTATAATCTTTTCAAAAAGCAAGCTA 1194

London  GluCl2 -----------------------------------AATACTCAAATAAAAATATATTTT 1222
SOL-BEi GluCl2 -----------------------------------AATACTCAAATAAAAATATATTTT 1222
MAR-ABi GluCl2 GAAACTTTGAACAAATTGTCACAATAAGAACAGGCCAAGACCGTGCTAATATTAACATTA 1254

London  GluCl2 CTCGCTG---GTGGTCAAAGTTTCCA---ACCCGATCTAAACGAATCGATGTTGTGTCGA 1276
SOL-BEi GluCl2 CTCGCTG---GTGGTCAAAGTTTCCA---ACCCGATCTAAACGAATCGATGTTGTGTCGA 1276
MAR-ABi GluCl2 ACCCATGATAATGATCGTACTCATCAGTTACTTGATAAGAAATCATCGGT---------- 1304

London  GluCl2 GAATATTTTTCCCGTTAATGTTTTGCTTATTCAATCTTGTC------TATTGGGTTACTT 1330
SOL-BEi GluCl2 GAATATTTTTCCCGTTAATGTTTTGCTTATTCAATCTTGTC------TATTGGGTTACTT 1330
MAR-ABi GluCl2 ---TAAATTTTCTTGTAATGATACGCTAAACTGTTCTTGGTGAGAAATATTGTTATAATA 1361

London  GluCl2 AT---------------------------------------------------------- 1332
SOL-BEi GluCl2 AT---------------------------------------------------------- 1332
MAR-ABi GluCl2 AGATCACAGTGAGAATGAAGAATACGAAATACGGATCCAAAGAATATTTGCAAAAACAAG 1421

London  GluCl2 ----------------------CTGTT------TCGCCATAAAAGAGATAAAAATGTTTA 1364
SOL-BEi GluCl2 ----------------------CTGTT------TCGCCATAAAAGAGATAAAAATGTTTA 1364
MAR-ABi GluCl2 AATTTGTCATTAATTTGACAATTTATTGCCAAGTTGGCAC---AAAGATGTAAATGATAT 1478

London  GluCl2 TTAA-------------------------------------------------------- 1368
SOL-BEi GluCl2 TTAA-------------------------------------------------------- 1368
MAR-ABi GluCl2 TTCATGGTAAGAATATTGAATTTCAATAGATTGACAAGATATTGAGAATAGAGGAATAAA 1538
```

**(b)**

```
London  GluCl2 VSLGVTTLLTMATQISGINASLPPVSYIKAIDVWTGVCLAFVFGALLEFALVNYASRSDA 360
SOL-BEi GluCl2 VSLGVTTLLTMATQISGINASLPPVSYIKAIDVWTGVCLAFVFGALLEFALVNYASRSDA 360
MAR-ABi GluCl2 VSLGVTTLLTMATQISGINASLPPVSYIKAIDVWTGVCLAFVFGALLEFALVNYASRSDA 360

London  GluCl2 HRHAREFNGLRYQRRWDRDGNVISDETSYALR----PL--------------VI---K- 397
SOL-BEi GluCl2 HRHAREFNGLRYQRRWDRDGNVISDETSYALR----PL--------------VI---K- 397
MAR-ABi GluCl2 HRHAREFNGLTRVILFSISWQAMTKRASLAIIFSKSKLETLNKLSQ*EQAKTVLILTLTH 419

               inserted site in MAR-ABi

London  GluCl2 -------------GSEYSNKNIFSRWWSKFPTRS-KR-----IDVVSRIFFPLMFC---- 434
SOL-BEi GluCl2 -------------GSEYSNKNIFSRWWSKFPTRS-KR-----IDVVSRIFFPLMFC---- 434
MAR-ABi GluCl2 DNDRTHQLLDKKSSVKFSCNDTLNCSW*EILL**DHSENEEYEIRIQRIFAKTRICH*FD 475

London  GluCl2 --------LFNLVYWVTYLFRHKRDKN-------VY*----------------------- 455
SOL-BEi GluCl2 --------LFNLVYWVTYLFRHKRDKN-------VY*----------------------- 455
MAR-ABi GluCl2 NLLPSWHKDVNDISW*EY*ISID*QDIENRGINRKYEIN*IRKLFFPGIFDYF*LIGQDL 530

London  GluCl2 ------------------------------------------------------------ 455
SOL-BEi GluCl2 ------------------------------------------------------------ 455
MAR-ABi GluCl2 GMVFNYPCWLMDGHGLGR*GGTQKNQQQENCEKLNLMNAVMMQQRTCQVLMALIRLTVMK 589

London  GluCl2 -------- 455
SOL-BEi GluCl2 -------- 455
MAR-ABi GluCl2 RLVLSIHH 597
```
