## Supplementary file 1 for "Intraspecific diversity in the mechanisms underlying abamectin resistance in a cosmopolitan pest"

>TuGluCl2_insertion sequence

ACCCGGGTAATACTTTTCAGCATTTCATGGCAAGCTATGACCAAACGAGCAAGCCTTGCTATAATCTTTTCAAAAAGCAAGCTAGAAACTTTGAACAAATTGTCACAATAAGAACAGGCCAAGACCGTGCTAATATTAACATTAACCCATGATAATGATCGTACTCATCAGTTACTTGATAAGAAATCATCGGTTAAATTTTCTTGTAATGATACGCTAAACTGTTCTTGGTGAGAAATATTGTTATAATAAGATCACAGTGAGAATGAAGAATACGAAATACGGATCCAAAGAATATTTGCAAAAACAAGAATTTGTCATTAATTTGACAATTTATTGCCAAGTTGGCACAAAGATGTAAATGATATTTCATGGTAAGAATATTGAATTTCAATAGATTGACAAGATATTGAGAATAGAGGAATAAATCGTAAATATGAGATCAATTGAATTAGAAAATTGTTTTTTCCTGGAATTTTTGACTATTTTTGACTAATTGGGCAAGATCTTGGAATGGTCTTTAATTATCCTTGCTGGTTGATGGACGGTCACGGTCTTGGCAGGTAGGGAGGGACGCAAAAAAACCAGCAGCAAGAAAACTGCGAGAAATTAAACCTGATGAACGCAGTCATGATGCAACAAAGAACTTGCCAAGTTCTCATGGCTCTAATTCGTCTAACAGTCATGAAAAGCCTGGTTTTGTCGATTCATCACCAAAAAATGATTTGTCTGAACCGAATCAGGATGATATTGACAGTGATGTAACGATATCAGGATCTCATGAAGAAAATCAGGAACAAGGAGATTCTTCACCGATCAATCTTCAGTCATTTAAAGTAAATCAAGTTGGCGACCTAAATGATAATCATTTTGAAGACGAAGAAACAATTGAATGATCTTGGCAATTTCTTGGCAAAATATTACTTATTATTTTTTGATACTTTTGAGTTTATATAGATTAAATTATTAAGAAATCTTTCAATATTTCAATAAAATTTTTTTGCAAATAATGACAAGATCTTATTTAGTTGTTGCCAAGAAATCAATCGTGAATCAGTGCAAAATCATGAAAAGATTGTGCACAAGAATACATCATGATTTTTAGCGAAAAATTAGCTGTTTCTTGCTGGTATTTCCTTTCATTTCTTGTGATGATTCGATTTTGTCTTTTGCCAAGGAAACAGTGAAGTCGTATCTATTTCTTTTTGAATTTTTCTGGTAAGATCAAACACTTATTAGTAGTGTTCCAGAAAATTTCTGACAATTTTTTGCCATGAAAGCATAAGAGAATCGACCGAAATTTTATCTTCTTAACTTGCTAATTCTTCAGTAATGCTGTGCCCTTCAAGAATCTTATATGACTTAATATGATCATATCAATAAAATATGCAGATAAATAGTCTGAACATGAGTAAAAAATTGTTAATGCATAACAAAAGCTTGATGGAAATACTCGACATTAAAATGTGCCAAGCGATTTGCTATATTCTTGTCGAAAAGTTCTTGCCCATTTTCATGCTATGACTGCCTGAAAGTATTACCCGGGA
